# An *HSP90-released reduced-eye phenotype alters light-dependent behaviour in Tribolium castaneum*

**DOI:** 10.64898/2026.04.02.716055

**Authors:** Tobias Prüser, R Reshma, Angelica Coculla, Ralf Stanewsky, Joachim Kurtz, Nora K. E. Schulz

**Affiliations:** Institute for Evolution and Biodiversity, University of Münster, Hüfferstr. 1, 48149 Münster, Germany; Institute of Neuro- and Behavioural Biology, University of Münster, 48149 Münster, Germany; Joint Institute for Individualisation in a Changing Environment, University of Münster and Bielefeld University; Department of Evolutionary Biology, Bielefeld University, 33615, Bielefeld, Germany

**Keywords:** evolutionary capacitance, HSP90, reduced-eye phenotype, light-dependent behaviour, adaptive benefits, *Tribolium castaneum*

## Abstract

Heat Shock Protein 90 (HSP90) functions as an evolutionary capacitor, allowing populations to store cryptic genetic variation that can be released under stress. While former studies have described the release of morphological variation, its behavioural consequences remain unexplored. In the red flour beetle, *Tribolium castaneum*, HSP90 inhibition released a phenotype with much smaller, less defined eyes that confers fitness benefits in continuous light and was subsequently assimilated. We hypothesized that altered eye morphology affects light perception and thereby changes light-dependent behaviours. To test whether phenotypes released via evolutionary capacitance can beneficially alter behaviour, we examined locomotor activity rhythm entrainment to light-dark cycles as well as individual and group light choice behaviour. Males of the reduced-eye phenotype exhibited a diminished startle response to sudden light exposure in locomotor activity assays. We also found reduced negative phototaxis in groups of beetles with reduced eyes. This modified behaviour, indicating reduced light sensitivity, may stem from impaired light perception caused by altered eye morphology. Lower light sensitivity could be beneficial under stressful environmental conditions by promoting the exploration of alternative niches. Therefore, this study provides the first evidence for potentially beneficial behavioural changes in a HSP90-released phenotype, reinforcing HSP90’s role as an evolutionary capacitor.

## Introduction

While natural selection acts at the phenotypic level, the raw material for evolutionary adaptation is provided by genetic variation arising from mutations. However, mutations occur at a low rate, are random and thus often deleterious, and may be removed by purifying selection in certain environments. Thus, the question arises, if there are other mechanisms allowing the maintenance of sufficient genetic variation. The evolutionary capacitor hypothesis proposes that certain buffering mechanisms allow populations to store some of their genetic variation in a cryptic form (cryptic genetic variation, CGV), without exhibiting its corresponding phenotype in a stable environment. But upon disturbance of the buffering system this CGV can be released as phenotypic variation [1]. Such mobilization of hidden genetic variation is hypothesized to speed up the process of adaptive evolution [2,3]. Heat shock protein 90 (HSP90) is the so far best-studied and most likely candidate to provide such evolutionary capacitance [1,4]. This chaperone is highly abundant in eukaryotes, involved in protein homeostasis [5] and appears to play an important role in the suppression of CGV [2,4]. In a situation of environmental stress such as heat shock, the increase in damaged and partially denatured proteins limits the availability of HSP90 [2]. Thus, the buffering function becomes depleted and the formerly CGV is expressed as phenotypic variation, on which natural selection can act [4,6,7]. Additionally, HSP90 is involved in the epigenetic regulation of gene expression through chromatin remodeling [8] and suppression of mutagenic transposons [9,10], providing further mechanisms to increase variation and thereby enhancing the adaptability of a population.

While most reports of HSP90-buffered phenotypes have focused on a potential capacity for morphological evolution, impairment of chaperone function has also increased variation in circadian locomotor activity behaviour in model species such as *Drosophila melanogaster* [11,12] and, although much more difficult to evaluate, in *Tribolium castaneum* [13]. Consequently, HSP90 additionally can act as a capacitor of behavioural variation [11–13]. However, this distinction between morphological and behavioural evolution ignores the intimate connection between these two aspects of an individual [14]. For instance, a considerable number of reported HSP90 buffered phenotypes displayed altered eye morphology [4,7,15], which in return can affect fitness foremost through triggered physiological and behavioural responses [16]. This highlights the importance to take downstream behavioural consequences of morphological changes into account.

Intriguing findings regarding evolutionary capacitance have emerged from studies using the model organism *Tribolium castaneum*. HSP90 was downregulated in naïve beetles upon cohabitation with wounded conspecifics, a situation where mobilisation of genetic variation could be beneficial for producing phenotypes relevant for immune responses [17]. Impairment of HSP90 through chemical inhibition as well as RNA interference led to the release of an HSP90 mediated eye phenotype in this beetle, which was stably inherited across offspring generations without the need of further impairment [15]. Affected beetles showed reduced eyes of approximately half the size of the wild-type eye, with a reduced number and poorly defined ommatidia. This reduced-eye phenotype demonstrated a higher fecundity under continuous light conditions [15]. Given the significant impact of the number of ommatidia and the size of their facets on the sensitivity and resolution of compound eyes [18], it seems likely that this phenotype directly affects the vision of the beetles and thereby might also lead to changes in light-dependent behaviours.

Here, we made use of this reduced-eye phenotype to address the often-overlooked question of consequences for behavioural traits of HSP90 buffered morphological phenotypes. As the reduced-eye beetles have a fitness advantage in continuous light compared to normal-eye conspecifics [15], we focused on light responses as one of the main behaviours mediated by the eyes. We compared phototaxis and light-synchronized diurnal rhythms of locomotor activity as two of the most archetypical light-dependent behaviours [19]. *T. castaneum* lives in stored grain products such as flour, burying into its food substrate and thus shows adaptations to a cryptozoic lifestyle, e.g. an increase in odorant receptors [20,21]. In coherence with this life style, it exhibits negative phototaxis [22]. However, findings supporting negative phototaxis in *T. castaneum* are not entirely consistent [23] and other studies report contradicting observations regarding the attraction of the beetles to differently coloured LED light, e.g. high versus no attraction to UV light [24,25]. Also, light stimuli can startle the beetle, as it responds with heightened activity [26], indicating stress upon light exposure. Despite living in dark enviroments, *T. castaneum* shows clear diel patterns in its activity on the population level [26], although with strong individual variation and a large proportion of arrhythmic beetles [26–28].

We hypothesized that the smaller eyes of the reduced-eye phenotype would diminish the light sensitivity of the beetles, as inter- and intraspecific comparisons in insects revealed a positive relationship of eye size and light sensitivity [29,30]. We therefore assumed to observe a reduction of the negative phototaxis in beetles with reduced eyes, as a consequence of their presumably lower light sensitivity. In addition, due to the lack of the *Drosophila melanogaster* type Cryptochrome (Cry) photoreceptor, it is most likely that the compound eyes are the predominant organ mediating light entrainment of diel activity patterns in *T. castaneum* [31]. Therefore, we hypothesized that the reduced eyes would alter the synchronization of daily activity patterns. Specifically, we proposed that this phenotype might result in limited entrainment under low light conditions or a weaker behavioural response to light changes.

## Materials and Methods

### Beetle strains and rearing

We used offspring generations from experimental lines of *T. castaneum*, derived from an HSP90-inhibited parental generation of our general Cro1 lab strain, which originated in Croatia in 2010 [15,32]. The HSP90-inhibition in the original parental generation was done via feeding the geldanamycin analogue, 17-Dimethylaminoethylamino-17-demethoxygeldanamycin (17-DMAG). Individuals of the F1 generation with normal eyes were crossed to establish a polymorphic line, with around 10 % of individuals stably exhibiting the reduced-eye trait. This proportion of reduced-eyed beetles remained stable across all subsequent generations [15]. The beetles of these experimental lines were maintained for seven years in 450 mL glass jars with approximately 100 g of a flour-yeast mixture, consisting of 5% brewer’s yeast and 95% organic wheat flour (Bio Weizenmehl Typ 550, dm-Drogeriemarkt), at 30°C, 70% relative humidity (RH) and 12h:12h light: dark cycles (LD).

To produce animals for the behavioural experiments, approximately seven-day old adults were allowed to mate and oviposit for 24 hours, on fresh flour. We later separated the pupae by sex and individualised them in 96-well micro reaction plates (type F, Sarstedt) on 0.08 g flour diet. After eclosion, we determined the eye phenotype of the beetles under a microscope and returned the beetles to the same 96-well plates.

### Locomotor activity measurement

Since small age differences may influence the behaviour of *T. castaneum* during the first days of the adult stage [33,34], all measurements were started in the third week after eclosion. The locomotor activity of reduced-eye and normal-eye beetles was monitored for seven days in standard 12 hours: 12 hours light-dark cycles (LD) and seven days in constant darkness (DD) using the Drosophila activity monitoring system (DAM5H-4, Trikinetics Inc., Waltham, MA, USA; referred to as “DAM system”, for details see [26]) in 70%RH, 30 °C (Fig 1a). We conducted the experiment using different light intensities of 20 lux (n=24), 100 lux (n=32 and n = 24), and 2000 lux (n=32 and n = 24), to test the interactions of light intensity with behavioural effects of reduced eyes and to ensure consistency of the observed effects. Average activity patterns of both phenotypes were plotted as histograms using the Flytoolbox in MATLAB [35,36], and the rhythmicity of individual beetles was determined using individual actograms in combination with autocorrelation curves [35] and χ^2^ periodograms (for details see[26]). We further quantified the increase of activity in response to the onset of light for each individual as a startle response index (SRI). This SRI was defined as the average activity in the 30 minutes after light onset divided by the average activity two hours before until two hours after this 30-minute window.

**Fig 1.**
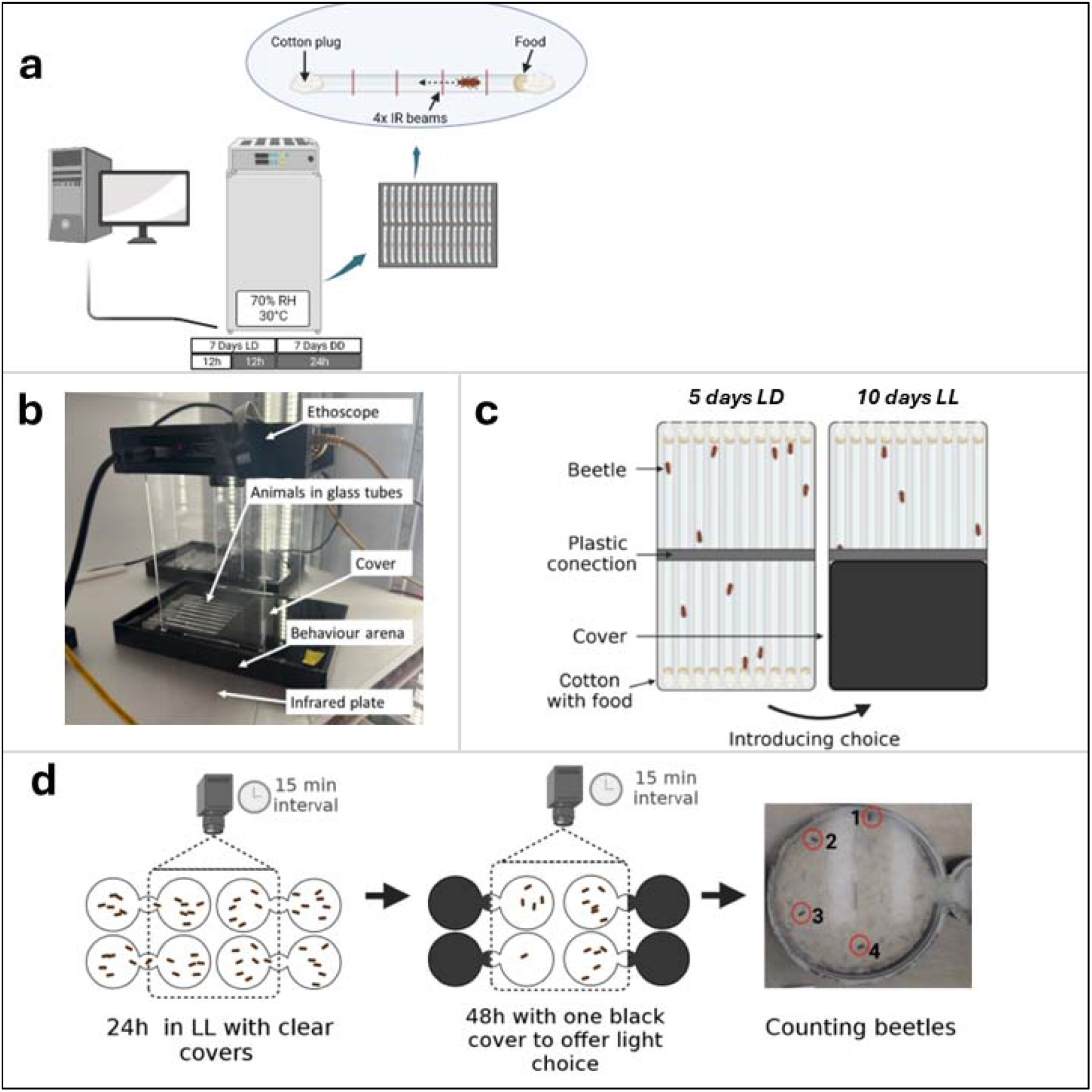
Overview of experimental procedures. **a** Locomotor activity assay. Individual beetles are placed in glass tubes, which are inserted into DAM monitoring systems and maintained in an incubator. Activity is quantified by recording each interruption of four infrared light beams that pass through the tube. **b** Photo of the Ethoscope mounted on the arena used for the light choice assay of isolated individuals. The arena with the Ethoscope is then placed on an infrared LED plate so that the infrared camera can track the movement of the beetles [37]. **c** Setup used for light choice assay of isolated individuals: The behavioural arena provides space for ten times two interconnected glass tubes, and by this for ten beetles. To divide side preference from light avoidance, we initially measured beetles for 5 days without cover in LD cycles. To initiate the 10 day choice phase of this experiment, one half of the tubes is covered so that the beetles can choose whether to stay in the light or in the dark. **d** Group light choice assay: Beetle groups were set into Carolina® choice chambers, together with food, and measured for 24 h in constant light without a cover, by taking pictures in 15 min intervals of one side of the arena. A dark cover was placed on the unmonitored side, and activity was recorded for 48 h during the choice phase. Photographs were analyzed by quantifying the number of beetles visible in each image.

### Light choice assay of isolated individuals

Individual light choice assays were performed using a choice apparatus adapted from the Ethoscope [38], a video tracking system for animals, consisting of a high-resolution infrared camera, connected to a small single-board computer (for details on adapted devices see [37]). The Ethoscopes were assembled in small plastic boxes (10×13×19cm), which were then placed over behavioural arenas to collect positional data of the beetles in the arena every ten seconds. For each individual, two interconnected glass tubes (*5mm* × *65mm*) were provided. Every glass tube was sealed on the non-connected side with cotton and contained a flour diet disc as food. Each behavioural arena can hold a total of ten glass tube pairs and therefore a total of 10 beetles (Fig 1b).

In this setup, we measured two-week-old male beetles of both phenotypes (n = 25 per phenotype). After assembling the Ethoscopes, the arenas were kept in an incubator (70% RH, 30 °C). The experiment consisted of two phases: in the first ‘no choice’ phase of the experiment, the beetles’ behaviour was measured in 12:12h LD cycles at 1000 lux for 5 days. In the second ‘choice’ phase for the following 10 days, the light condition was changed to constant light (LL-1000 lux) and a cover, made from infrared permeable material, was added to one side of the set-up. Hereby, the arena was divided into equally sized, constantly dark and constantly illuminated areas, providing a light choice to the beetles. One of the five arenas remained uncovered to serve as a control ensuring that changes in side preference during the second phase of the experiment were caused by the added cover. To assess the beetles’ response to light and distinguish it from random side bias, we first–during the no-choice phase of the experiment–measured the time the beetles spend on the side of the arena, which would be covered during the second phase, the choice phase of the experiment. To calculate the light choice of each individual, we then subtracted the percentage of time it spent on the newly covered side of the arena during the choice phase from the time spent on this side during the no-choice phase (light choice = time (%)_no choice phase_ – time (%)_choice phase_).

### Light choice assay in groups

To study the light choice of beetles in a group-setting, we transferred individualised beetles, seven days after eclosion, to petri dishes containing 5g of flour. Each petri dish housed a group of 20 beetles (10 males and 10 females). Groups consisted homogenously of either wildtype or reduced-eye beetles and were kept under these conditions for seven days for acclimatisation to the social environment. The groups were then transferred into choice chambers (21,59 x 1,9 x 9 cm, Carolina Biological Supply Co), with edges coloured black (Loopcolors Germany-LP-104 Black). These chambers served as the behavioural arenas. The surface of the arenas was covered with a thin layer of flour. Arenas were placed into an incubator (70% RH, 30°C) with constant light (100 lux). Starting on the second day, a camera mounted above the arenas (Logitech StreamCam) took pictures of one side of each arena in 15-minute intervals. During the first 24 hours of the experiment both sides of the arena were covered with a clear lid to test for side preferences. The following 48h of the experiment served to measure phototaxis. For this, a black cover was added to the unmonitored side of the arena (Fig 1c). The experiment was replicated three times with four arenas (two per phenotype) under the same conditions. The captured images were analysed by counting the number of visible beetles in each image. Two people counted either half of the pictures, while being blind to the treatment groups. Repeatability of counts was confirmed by counting beetles in the same subsample of 100 images (Spearman correlation coefficient between assessor counts = 0.966). Beetles that were not clearly identifiable or on the dividing line were counted as half. It should be noted that we counted on average only 7 out of 20 beetles in the no-choice phase, while 10 would be expected in case of a random distribution of the beetles. This underestimation was likely due to beetles that remained undetected because they buried into the flour or because mating beetles were counted as one single beetle. Importantly, this effect was consistent for both phenotypes and thus does not bias our results.

For analysis, the average number of beetles counted was calculated for each replicate and day. The average number of beetles counted between the first day of ‘no choice’ and the following days of ‘choice’ was then compared for both phenotypes to determine whether the beetles avoid the light, when given the opportunity.

### Statistical analysis

All statistical analyses were carried out in R [39] operated with the interface of RStudio [40]. We applied Generalized Linear Mixed Models using the package glmmTMB [41] after determining the error distribution that best represented the data from each experiment. Random factors were chosen in a manner that they accounted for repeated measurements to avoid pseudo-replication and to control for block effects. Model details are given in Table 1. The data was visualised using the standard ggplot2 [42] toolkit, with additional implementations of Patchwork [43].

**Table 1.** overview of GLMMs used for statistical analysis.

|  | Response variable | Error distribution | Fixed factors | Random factors |
| --- | --- | --- | --- | --- |
| *Frequency of rhythmic behaviour* | rhythmicity | binomial | sex, light condition, phenotype | ID, Experiment light intensity |
| *Startle response* | startle response index | beta | sex, light intensity, phenotype | experimental block |
| *Individual light avoidance* | phototaxis | beta | phenotype, cover | position in ethoscope |
| *Group light avoidance* | proportion of counted beetles | beta_family (link=log) | day | replicate |

## Results

### Reduced eyes do not alter circadian behavioural rhythms but reduce light sensitivity in male beetles

As the compound eyes are assumed to be the principal circadian photoreceptor in beetles [31], we tested whether the reduced-eye phenotype affects the beetles’ ability to entrain their locomotor activity to light-dark cycles (LD). We monitored the locomotor activity of both phenotypes from the polymorphic lines using the DAM system in LD (L = 20 lux, 100 lux and 2000 lux) and DD and compared the average locomotor activity patterns and the proportion of rhythmic beetles (Fig 2 & 3). Average histograms of both phenotypes showed comparable activity patterns (Fig 3). Both the reduced-eye (RE) and normal-eye (NE) beetles exhibited sex- and light-dependent differences in their behaviour. In both phenotypes across all light intensities, most males exhibited rhythmic behaviour in LD (males NE: 82.8%; RE: 69.4%, Fig 2, Table S1). The proportion of rhythmic beetles significantly decreased by more than half in DD (GLMM: p<0.001, males NE: 32.84%; RE:26.12%, Fig 2; Table S1, Table S2). The number of rhythmic females was significantly lower than the number of rhythmic males under LD (GLMM p<0.001, females NE: 43.4%; RE: 42.1%, Fig 2, Table S1, Table S2) and DD (females NE: 19.1%; RE: 18.8%, Table S1) but the difference between LD and DD was less distinct compared to males (GLMM: p=0.002 Table S2). While there was distinctly no difference between the two phenotypes in females (GLMM: p=0.82, Fig 2, Table S2), there was a trend towards less rhythmic individuals in males with reduced eyes (p=0.083, Table S2, Fig 2).

**Fig 2.**
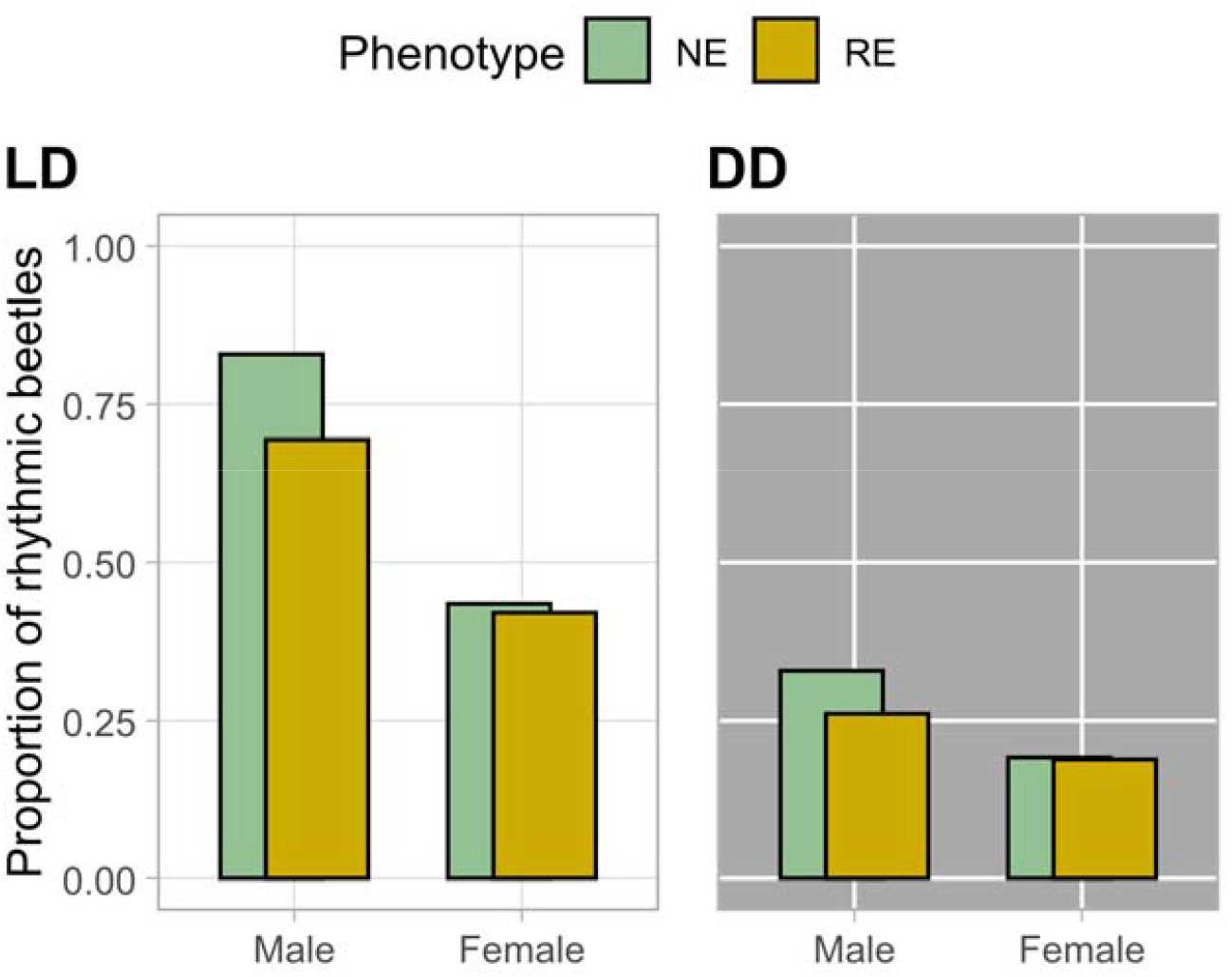
Proportion of normal (NE) and reduced-eye (RE) rhythmic beetles under LD and DD. Bar plot showing the proportion of rhythmic beetles under LD and DD for all experiments (NE: males n=134, females n=136, RE: males n= 133, females n= 134; data combined for all light intensities see Table S1). The rhythmicity of each beetle was determined by the evaluation of individual actograms in combination with autocorrelation functions and χ2 –periodograms for both light conditions.

**Fig 3.**
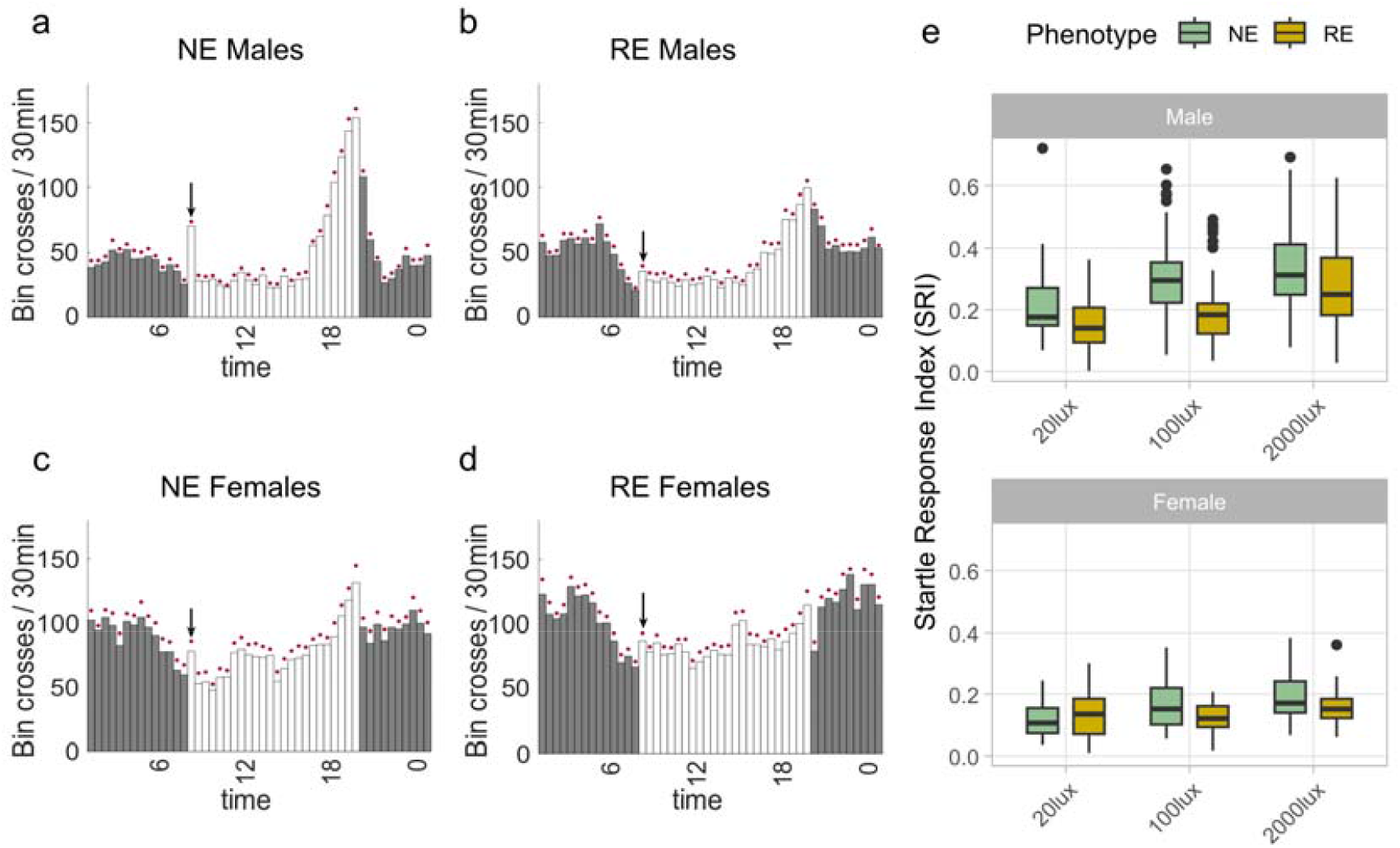
Reduction of startle response in reduced-eye males. **a-d** Average histograms of normal eye (NE) and reduced-eye (RE) males and females showing the average activity patterns for all beetles in a group during the 7 days under LD (L=100 lux) in 30-minute bins. Light is indicated by shading (white=lights on, grey=lights off), (NE males n=31, RE males n=31, RE females n=30, NE females n=32). For average histograms corresponding to other light intensities, see Fig S1. The 30-minute interval counted as a startle response (immediately following the turning on of the light) is indicated by an arrow above the bin. e Box plots showing startle response index (defined as the average activity of a beetle during 30min after the light turns on, normalized by its activity in a 4.5-hour period starting from two hours before until two hours after) of beetles compared between light intensities, phenotypes and sexes. For sample sizes see S1.

However, males of the two phenotypes showed a notable difference in startle response (strongly heightened activity in the 30min after the onset of light in the morning; Fig 3a-d, arrow). There was a significantly reduced reaction, as measured by the startle response index (SRI) described above, in reduced-eye males (GLMM: p=0.029, Fig 3b, Table S3) irrespective of the light intensity (Table S3). Thus, the reduced eyes seem to alter the light response of the beetles, which may indicate an altered light reception of this phenotype. We also observed a lower startle response for reduced-eye females compared to the normal-eye phenotype in 100 lux and 2000 lux light condition (Fig 3e, Fig S1, Table S3), but this reduction was not statistically significant in either condition. It seems therefore possible that the reduction effect is also occurring in the females but is not detected by the model because of the overall less pronounced startle response in this sex.

### Reduction in negative phototaxis in groups of reduced-eye beetles

Since we found an indication of altered light reception for reduced-eye beetles, we further investigated other behavioural consequences of this altered light response. Thus, we measured phototaxis of individual beetles using the Ethoscope, a video tracking system by measuring the proportion of time the beetles spend in the dark versus light side of a glass tube. We account for any light condition independent side bias by normalizing data against an initial measurement without a covered area. Both phenotypes showed negative phototaxis, defined here as the differences in side preference before and after covering the behavioural arenas (mean differences for beetles with normal eyes: 14.95%, for beetles with reduced eyes: 15.44%). This was confirmed by comparison with the uncovered control group, which showed no corresponding change in side preference (p-value=0.051, Fig S2, Table S4). However, there was no difference between the two phenotypes (GLMM: p=0.825, Fig 4a, Table S4).

**Fig 4.**
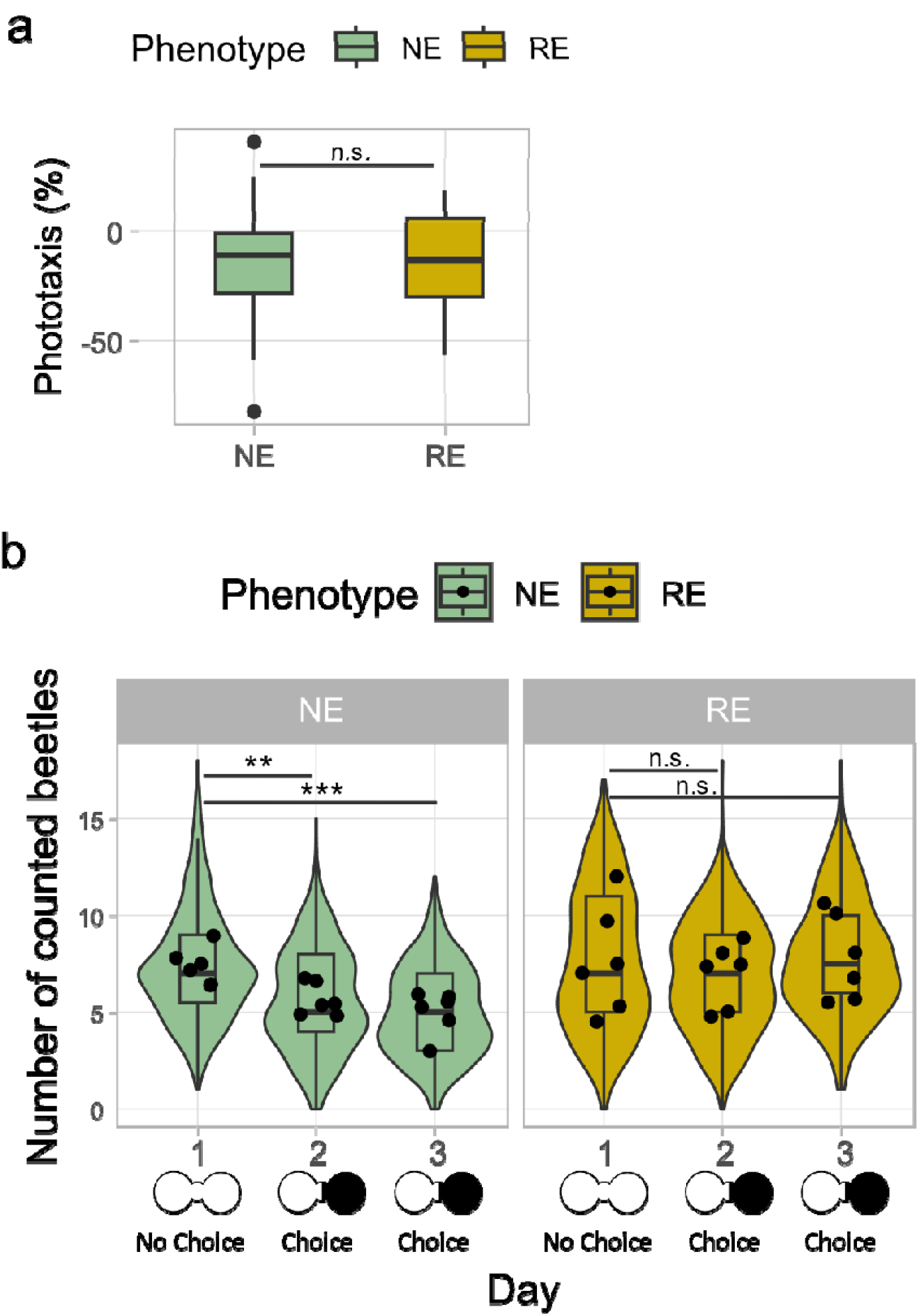
Phototactic response of reduced eye and normal eye beetles in isolation and groups. **a Negative phototaxis of reduced-eye and normal eye beetles in isolation**. The phototaxis was defined as the differences in side preference (light vs dark) before and after behavioural arenas were covered (RE: n=19, NE: n=18). Positive values display preference of the illuminated side, negative values indicate light avoidance. **b Reduction of negative phototaxis of reduced-eye beetles in groups**. Shown is the number of beetles counted on the monitored side (out of a total of 20 beetles per arena). The violin plot shows the variation in the number of beetles counted across all replicates and time points per day, separated by phenotype, which is also summarized in the overlayed boxplot (96 measurements per replicate per day, 6 replicates per phenotype). Sketches below the arena indicate the status of the arena (uncovered on the first day - no choice, covered on one side on the second and third day - choice). The mean values of the arena used for the statistical analysis are shown as black dots in the diagrams

As light choice in *T. castaneum* is influenced by interactions with conspecifics [22], we additionally measured the light choice behaviour for the phenotypes in a group setting using choice arenas. We counted the number of beetles in one half of the arena for 24 h in constant light, and afterwards for further 48 h, but with a cover added to the non-counted side of the arena. On the first day, when there was no covered dark space offered, on average 7.7 (out of 20) beetles per arena and time point were counted on the monitored side in reduced-eye groups and 7.4 (out of 20) in the normal eye groups. After the beetles were given a choice to avoid the light, the number of normal eye beetles on the light side decreased significantly to an average of 5.7 beetles (GLMM p<0.01, Fig 4b, Table S5) on day two, and further to 5.0 on the third day (GLMM p<0.001, Fig 4b, Table S5), indicating negative phototaxis. Contrary to this, there was no significant light avoidance in reduced-eye groups, neither during the second day (average number of beetles =6.9, p= 0.467, Table S6) nor during the third day of the experiment (average number of beetles = 7.8, p=0.871, Table S.6) providing evidence for an altered group phototaxis in the reduced-eye phenotype.

## Discussion

HSP90 is the most extensively studied and characterised evolutionary capacitor [44], known to buffer morphological variation in a range of animal models [4,7,15,45]. However, whether these altered morphologies have secondary effects on the behaviour has so far remained unexplored. For example, it is often assumed that reduced eyes in cryptozoic species are adaptive, primarly because of energy conservation [46]. But what about subsequent changes in behaviour, brought about by reduced light perception? Can we observe such changes and could they be beneficial?

To address these questions, we here made use of a recently characterised HSP90-regulated phenotype in *T. castaneum*. In the present study, reduced-eye beetles entrained to light–dark cycles, with only a slight reduction in rhythmically behaving males, and showed daily activity patterns largely similar to those of normal-eyed beetles. For this, it could be relevant that eyes are only reduced in size and not completely absent and therefore most likely remain functional as photoreceptors. In *D. melanogaster*, even the total absence of compound eyes does not impede the fly’s ability to entrain to light-dark cycles, because of its Cry-dependent extraretinal photoreception [47]. There is also evidence for extraretinal photoreception in beetles [48] and it is so far not known whether these structures differ in the reduced-eye phenotype.

While reduced-eye beetles were able to entrain normally to light-dark cycles, we observed a reduction in the startle response in reduced-eye males when lights are switched on in the morning. The disappearance of such startle response in eyeless mutants of *D. melanogaster* [49] and its persistence in clock-disturbed flies [50] indicate that these activity peaks are eye-mediated responses to environmental changes, rather than clock-controlled behavioural patterns. Similarly, in *T. castaneum*, this startle response is considered to be independent of the circadian clock [26]. The reduced-eye size may lead to a lower response to light stimuli due to reduced light sensitivity. This hypothesis is supported by reports in butterflies and bumblebees, in which larger individuals with larger eyes demonstrate more light-sensitive vision [29,30]. It should be noted that increased sensitivity in the described cases stems from larger facet size, whereas reduced-eye beetles mainly differ in facet number and shape [15], which likely also reduces light capture. In contrasts to males, the reduction of the startle response was not significant in females, which could be due to the fact that also normal-eye females exhibit a weaker startle response than males, i.e. potential differences between phenotypes are more difficult to spot in females. Sex-dependent differences in light response are known in other insect species, e.g. differences in retinal responsivity [51] or phototaxis[52].

To evaluate additional behavioural changes arising from the presumably altered light sensitivity of the reduced-eye beetles, we tested the light choice behaviour of both phenotypes. We found similar light avoidance for reduced- and normal-eye males in isolation under a high light intensity of 1000 lux. The group choice assays provided further evidence for a moderate negative phototaxis of normal-eye *T. castaneum* under a low light intensity (100 lux). The consistency of light avoidance of *T. castaneum* between experiments seems noteworthy itself, since reports on the phototaxis of this beetle are contradictory [22–25,53–56]. Besides differences in experimental design, the use of different *T. castaneum* strains could be a possible reason for the contradictory observations in this species. Other light-dependent behaviours, such as locomotor activity rhythms in light–dark cycles, also seem to differ between beetles collected from different geographical locations, with Cro1 beetles being mostly active during later times of day [26], whereas other populations’ activity peaks early in the day [57].

Furthermore, in contrast to conventional phototaxis experiments, we measured phototaxis not at a specific time point but as the proportion of time spent in the dark over several days. Animals actively shape the light environment that they experience in a time-of-day-dependent manner [58], and the sensitivity of compound eyes can change over the course of the day [59]. Thus, averaging behaviour over several days may provide a clearer picture of overall behaviour than measurements at specific time points.

Most interestingly, in the group measurements, the negative phototaxis was absent in the reduced-eye beetles, providing evidence for an altered light choice of this HSP90-mediated phenotype. The correlation between smaller eyes and weaker phototaxis has been observed in both intra- and interspecific comparisons. For instance, larger bumblebees with accordingly bigger eyes exhibit stronger positive phototaxis at low light intensities [60]. Similarly, the cave beetle *Ptomaphagus hirtus*, with significantly smaller eyes than its relative *P. cavernicola*, shows distinct but weaker negative phototaxis despite a similar ecology [23]. However, attributing these differences to eye size alone remains challenging. In bumblebees, body size is associated with division of labour and in comparisons across species it cannot be ruled out that the altered phototaxis is not solely due to sensory limitations but reflects adaptive shifts in phototactic preferences between species. In the present study, however, we directly compare reduced-eye beetles with their normal-eyed relatives from the same genetic background, making it likely that observed differences stem from eye morphology—though we cannot fully exclude other unrecognized phenotypic changes.

The contrasting results between the individual and the group choice assay likely stem from the variation in light intensity (100 lux group choice vs 1000 lux individual choice), as strength of illumination influences phototaxis [60,61] and therefore differences sometimes only become apparent under lower light intensities [60]. Higher light intensity was used in individual tests because even normal-eye beetles did not display negative phototaxis at lower intensities in preliminary tests in Ethoscope experiments. This might be caused by the small size of the measurement tubes, as factors such as size of the arena can alter the behaviour of *T. castaneum* [62]. At low light intensity the intrinsic locomotor activity of the beetles possibly outweighed their light avoidance and led to movement throughout both sides of the tubes.

In addition to differences in light intensity, group measurements more closely resembled the beetle’s natural environment [63], and may have additionally influenced the outcome. For example, in group living zebra finches negative effects of artificial light at night seem to be stronger in groups, presumably driven by extremely sensitive individuals [64]. Importantly, social cues can also modify individual responses to light in insects, as for example in honey bees, where they can overwrite photic entrainment of circadian rhythms [65] Finally, the social environment such as the groups’ sex composition can affect phototaxis in *T*.*castaneum* [22].

The hypothesis of evolutionary capacitance proposes that the release of CGV may occasionally enhance an organism’s phenotype-environment match, albeit most released variation is supposed to be detrimental [4,6,66]. The reduced-eye phenotype could provide a rare example for fitness gains under certain conditions, as reduced-eye beetles were found to produce more offspring than normal eye beetles in continuous light [15]. The present study potentially provides a behavioural explanation for this observation: the reduced startle response in reduced-eye males might indicate that the individuals experience less stress as a consequence of their reduced light sensitivity. Reduction in eye size as an adjustment to stressfull light conditions has been reported in *Daphnia magna*, which react plastically with smaller eyes to stressfull UV radiation [67]. Our findings further indicate that the diminished light sensitivity has resulted in a decline in negative phototaxis in beetle groups of the reduced-eye phenotype. The evolutionary ecology of phototactic behaviours is diverse, complex and often remains unclear [68]. Nevertheless there are striking examples of strong ecological relevance such as parasites manipulating phototaxis to increase transmission success [69,70]. For dark-dwelling animals, such as subterranean amphipods or cave-dwelling spiders, phototaxis is proposed to be a habitat choice mechanism, i.e. a form of niche choice [71], ensuring that the species avoids leaving the environment to which it is best adapted [72,73]. However, in a situation, where the organism is facing stress in its natural environment, it may be beneficial if a lowered light avoidance makes it less reluctant to leave its habitat and expand its ecological niche. This might also lead to enhanced dispersal of the species. In line with this argument, the evolutionary capacitor hypothesis predicts that CGV such as the reduced-eye phenotype is released in stressful environments to enable rapid adaptation. Taken together we provide further evidence of how an HSP90-released phenotype could help improve the fitness of an organism under certain environmental conditions by altering its behaviour.

To our knowledge, this is the first demonstration of potentially adaptive behavioural consequences of formerly cryptic genetic variation. Together with the direct fitness benefits of the reduced-eye phenotype [15], this fortifies the hypothesis of HSP90 as an evolutionary capacitor and its role in accelerating the process of evolutionary adaptation.

## Acknowledgement

We would like to thank Maite Ogueta for valuable input on analyses. Further we would like to thank Luis Garcia-Rodriguez, who built the Ethoscopes and developed the choice behavioural arenas used with them.

## Author contributions

T.P.: conceptualization, data curation, formal analysis, investigation, methodology, project administration, visualizaiton, writing-original draft, writing-review & editing, R.R.: conceptualization, data curation, formal analysis, investigation, methodology, project administration, visualizaiton, writing-original draft, writing-review & editing, A.C.: formal analysis, writing-review & editing R.S.: writing-review & editing, resources, J.K.: conceptualization, funding acquisition, writing-review & editing, resources, supervision, N.K.E.S.: conceptualization, formal analysis, methodology, project administration, writing-review & editing, supervision.

## Data accessibility

All data presented in this study, as well as all R code used for statistical analysis and to produce the figures will be made available upon reasonable request and will become publically available upon publication in a peer reviewed journal.

