## supplements for "An *HSP90-released reduced-eye phenotype alters light-dependent behaviour in Tribolium castaneum*"

**Table S1 Number of rhythmic beetles for each experiment**

|  |  | **Experiment** | | | | | |
| --- | --- | --- | --- | --- | --- | --- | --- |
| Sex | Phenotype | **Light condition** | 1  (20lux) | 2  (100lux) | 3  (2000lux) | 4  (100lux) | 4  (2000lux) |
| Male | **RE**  **RE** | **LD** | 15/23 | 24/31 | 21/31 | 18/24 | 15/24 |
|  |  | **DD** | 4/23 | 8/31 | 5/31 | 12/24 | 7/24 |
|  | **NE**  **NE** | **LD** | 17/23 | 24/31 | 29/32 | 22/24 | 19/24 |
|  |  | **DD** | 9/23 | 7/31 | 8/32 | 11/24 | 9/24 |
| Female | **RE**  **RE** | **LD** | 13/24 | 8/30 | 18/32 | 10/24 | 7/24 |
|  |  | **DD** | 4/24 | 4/30 | 7/32 | 3/24 | 7/24 |
|  | **NE**  **NE** | **LD** | 11/24 | 11/32 | 18/32 | 10/24 | 9/24 |
|  |  | **DD** | 7/24 | 6/32 | 6/32 | 3/24 | 4/24 |

| **Table S2\| Analysis of proportion of rhythmic beetles.** GLMM with binomial error distribution (Rythmicity ~ Sex * Phenotype*Light condition + (1 \| Experiment) + (1 \| ID) + (1 \| Intensity) | | | | |
| --- | --- | --- | --- | --- |
| **Fixed effects** | **Estimate** | **Std. Error** | **z value** | **p value** |
| Intercept | *-0.362* | *0.233* | *-1.557* | *0.112* |
| Sexmale | *2.419* | 0.394 | 6.133 | < 0.001 |
| PhenotypeRE | *0.075* | 0.330 | -0.227 | 0.820 |
| LightDD | *-1.531* | 0.330 | -4.361 | < 0.001 |
| Sexmale:PhenotypeRE | *-0.863* | 0.497 | -1.736 | 0.083 |
| Sexmale:LightDD | *-1.509* | 0.475 | -3..179 | 0.002 |
| PhenotypeRE:LightDD | *-0.026* | 0.453 | 0.057 | 0.954 |
| Sexmale:PhenotypeRE:LightDD | *0.493* | 0.650 | 0.761 | 0.447 |

| **Table S3\| Analysis of startle response**. GLMM with beta family error distribution (response ~ Phenotype*Sex*Light+(1\|Experiment)) | | | | |
| --- | --- | --- | --- | --- |
| **Fixed effects** | **Estimate** | **Std.Error** | **z value** | **p-value** |
| Intercept | *-1.903* | *0.204* | *-9.348* | < 0.001 |
| PhenotypeRE | *0.026* | 0.187 | 0.138 | 0.890 |
| Sexmale | *0.612* | 0.176 | 3.479 | <0.001 |
| Light100Lux | *0.436* | 0.238 | 1.837 | 0.066 |
| Light2000Lux | *0.372* | 0.237 | 1.570 | 0.117 |
| PhenotypeRE:Sexmale: | *-0.558* | 0.255 | -2.189 | 0.029 |
| PhenotypeRE:100Lux | *-0.341* | 0.221 | -1.546 | 0.122 |
| PhenotypeRE:2000Lux | *-0.246* | 0.217 | -1.134 | 0.257 |
| Sexmale:100lux | *0.095* | 0.205 | 0.466 | 0.641 |
| Sexmale2000lux | *0.118* | 0.203 | 0.581 | 0.561 |
| PhenotypeRE: Sexmale:Light100Lux | *0.340* | 0.299 | 1.136 | 0.256 |
| PhenotypeRE: Sexmale:Light2000Lux | *0.467* | 0.294 | 1.586 | 0.113 |

**Table S4** **Analysis of change in the side preference of isolated males of both phenotypes**.

| GLMM with beta family (link = "logit") error distribution (`Transformed data for Analysis`~ Cover*Phenotype +(1\|Position)) | | | | |
| --- | --- | --- | --- | --- |
| **Fixed effects** | **Estimate** | **Std. Error** | **z value** | **p-value** |
| Intercept | -0.337 | 0.125 | -2.704 | 0.007 |
| Non-covered group | 0.508 | 0.260 | 1.951 | 0.051 |
| PhenotypeRE | 0.038 | 0.172 | 0.221 | 0.825 |
| Non-covered:REPhenotype | -0.124 | 0.383 | -0.324 | 0.746 |

| **Table S5 Analysis of light avoidance of groups in NE phenotype** GLMM with beta family (link = "logit") error distribution (``Proportional_Counts ~ Day+ (1 \| `Replicate`)) | | | | |
| --- | --- | --- | --- | --- |
| **Fixed effects** | **Estimate** | **Std. Error** | **z value** | **p-value** |
| Intercept | *-0.535* | *0.085* | *-6.284* | <0.001 |
| Day2 | *-0.394* | 0.125 | -3.164 | 0.002 |
| Day3 | *-0.568* | 0.127 | -4.456 | <0.001 |

| **Table S6 Analysis of light avoidance of groups in RE phenotype** GLMM with beta family (link = "logit") error distribution (`Proportional_Counts` ~ Day+ (1 \| `Replicate`)) | | | | |
| --- | --- | --- | --- | --- |
| **Fixed effects** | **Estimate** | **Std. Error** | **z value** | **p-value** |
| Intercept | *-0.485* | *0.178* | *-2.721* | 0.007 |
| Day2 | *-0.151* | 0.208 | -0.727 | 0.467 |
| Day3 | *0.033* | 0.205 | 0.162 | 0.871 |

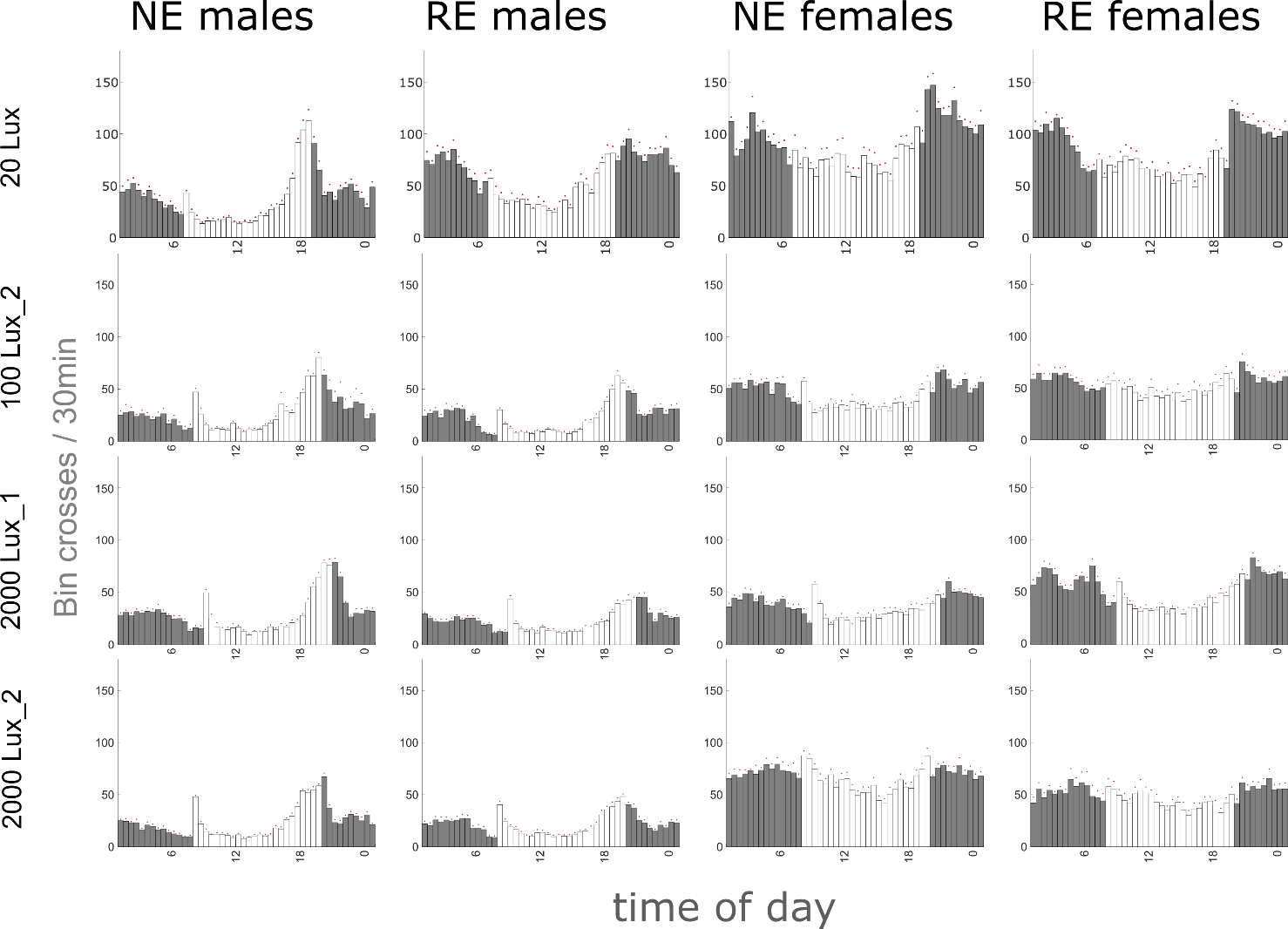
**Fig S1.** Average histograms of normal eye (NE) and reduced-eye (RE) males and females showing the average activity patterns for all beetles in a group during the 7 days under LD in 30-minute bins, separated by light intensity and experimental trial. Light is indicated by shading (white=lights on, grey=lights off). (20Lux: NE males n=23, RE males n=23, RE females n=24, NE females n=24 ; 100Lux_2 NE males n=24, RE males n=24, RE females n=24, NE females n=24 ; 2000Lux_1 : NE males n=32, RE males n=31, RE females n=32, NE females n=32 ; 2000Lux_2: NE males n=24, RE males n=24, RE females n=24, NE females n=24).

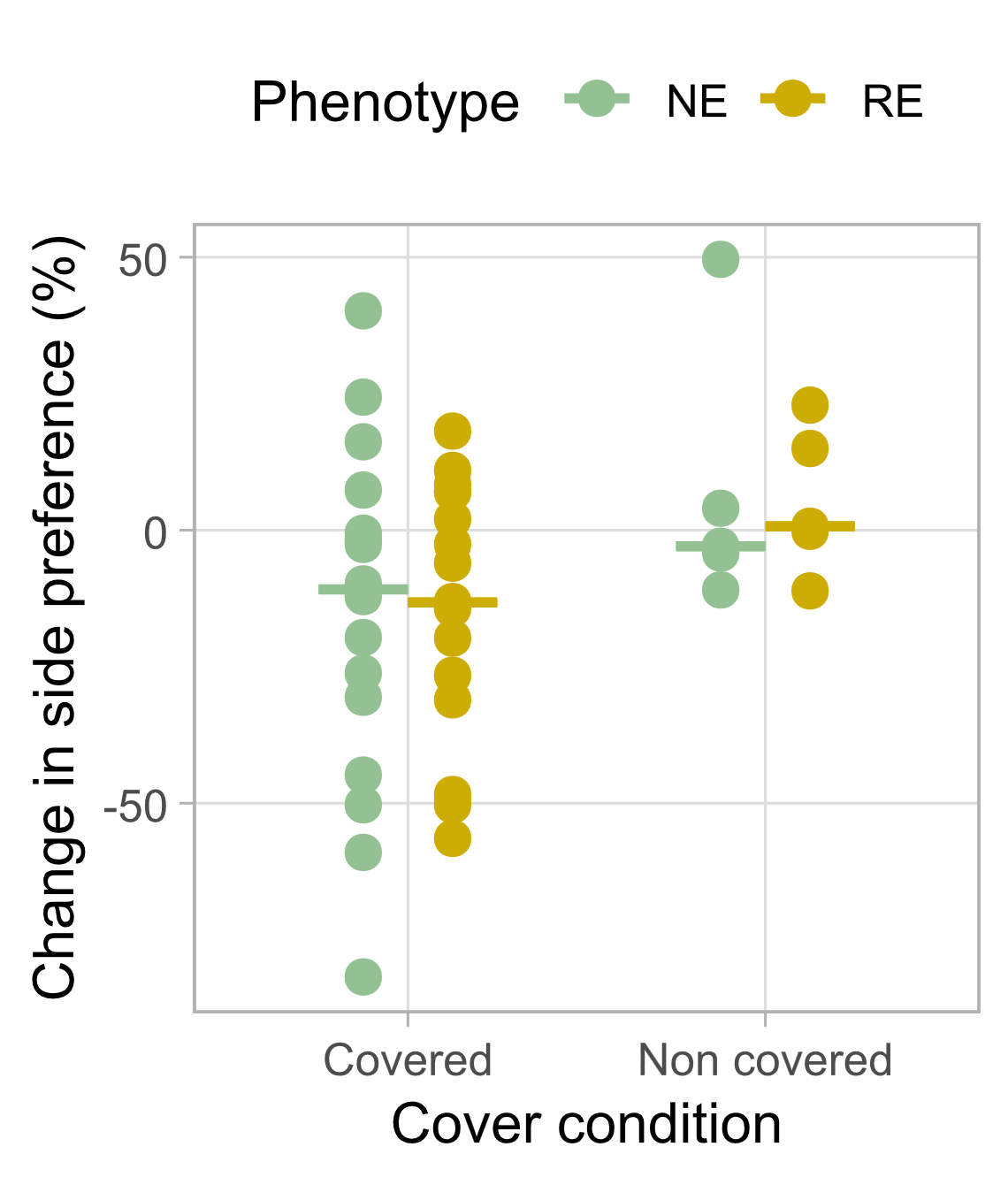

**Fig S2.** Comparison of the change in the side preference between RE and NE beetles after a cover was added to the "covered" treatment group. A non-covered arena served as control. (NE covered n=18, RE covered n=19, NE non covered n=5, RE non covered n=5).
